# Newly identified genomic sequences establish benchmarks for proposed taxonomic classification of jingmenviruses

**DOI:** 10.64898/2025.12.01.691605

**Authors:** Agathe M.G. Colmant, Rhys H. Parry, Rémi Charrel, Bruno Coutard

## Abstract

Jingmenviruses are a group of viruses related to orthoflaviviruses characterized by a segmented genome and multipartite organisation that have been detected worldwide in a wide range of hosts. As next-generation sequencing has become more affordable, increasing numbers of large-scale metagenomics studies have been published, alongside raw sequencing data. With the growing number of new jingmenvirus sequences identified in metagenomics data, it can be difficult to assess whether a new sequence is associated to a new virus species or to a strain of an existing species. In order to propose clear classification criteria for this group, we assembled a large jingmenvirus sequence database, from published and newly assembled sequences. Indeed, we screened data from studies that did not search for or report jingmenvirus sequences, looking for new strains of known jingmenvirus species. We then performed multiple sequence alignments and used the inferred percentage identity values to determine demarcation criteria based on the distribution of evolutionary distances upon pairwise comparisons. We report the identification of almost 60 libraries containing jingmenvirus sequences, in a wide range of sample types and geographical locations. Using these data and published jingmenvirus sequences, we have determined that to classify jingmenvirus sequences into virus species, at least four segments are required, on which eight cut-off values in percentage identity (nucleotide and amino acid) are used for demarcation. The ratification of this proposal would enhance consistency in virus taxonomy and provide a standardized framework for comparative genomics studies of jingmenviruses, a group that is yet under-characterised.

**Importance:** There are currently no guidelines to classify jingmenvirus sequences which means sequences are associated to species with no ratified rationale, which can result in missed opportunities at better describing jingmenvirus genomics. In this study we propose criteria to provide a parsimonious framework for classifying jingmenvirus species based on current knowledge, and allow us to propose re-classification of some sequences and to identify outlier sequences which we recommend should be further characterised *in vitro*.

## Introduction

Jingmenviruses are viruses with a segmented genome that have been detected all over the world in a wide range of samples, mostly by metagenomics (1). The two segments of the genome containing coding sequences for (i) the RdRp and methyltransferase domains as well as (ii) the helicase and protease domains share sequence homology with members of the *Orthoflavivirus* genus in the *Flaviviridae* family. The other segments contain coding sequences for putative structural proteins and share limited sequence homology with the rest of the virus realm, although structural homology has been found between some of these ORFs and the structure of orthoflavivirus structural proteins (2,3).

Jingmenviruses were not classified *per se* as of the ICTV August 2024 release and are referred to as unclassified viruses, related to the *Orthoflavivirus* genus, *Flaviviridae* family, *Amarillovirales* order in the NCBI and ICTV taxonomy browsers. Within the jingmenvirus group, no guidelines or criteria to associate sequences to viral species exist to date, which makes the classification of jingmenviruses challenging. In addition to this, the variability in identified segment numbers and the existence of jingmenvirus-related endogenous viral elements (EVE) integrated in tick genomes complicates annotation of novel sequences and therefore their classification (4–8).

To date, the published jingmenvirus sequences have been unofficially separated in two main clades based on phylogenetic analyses, one clade termed tick- and vertebrate-associated jingmenviruses and the other clade termed insect- and plant-associated jingmenviruses (1). While this designation facilitates sequence description, it is flawed in the sense that some sequences in the tick- and vertebrate-associated clade originated from mosquitoes, cattle feces or soil metagenomic samples while some sequences from the insect and plant-associated clade were derived from metagenomics crayfish, scorpions, water samples or even from vertebrates (1,4,9,10). No clear or official classification has been established yet.

Demarcation criteria to classify virus sequences can be determined using the distribution of percentage identity upon pairwise comparison of related nucleotide or amino acid sequences (11,12). The alignments from which these values are inferred ideally need to include sequences of different strains of the same virus as well as sequences of different virus species, in order to determine criteria that properly differentiate species from each other. In this study, we endeavour to propose criteria to clarify the taxonomic classification of jingmenviruses.

## Results

### New jingmenvirus sequences assembled from 59 libraries

In order to build a large jingmenvirus genomic sequence database, we searched for undescribed sequences of known jingmenviruses from both putative clades in public sequencing data. Using Serratus, we found 59 libraries with nucleic acid related to jingmenviruses that were not linked to a publication reporting jingmenvirus sequences and assembled full and partial jingmenvirus genomic sequences from these libraries. Of note, libraries SRR21218428, SRR14866274, SRR14866275, SRR14866276, SRR14866277 contain jingmenvirus sequences that have been partially described in publications but that were not deposited on Genbank (8,13).

Overall, we report 15 new sequences closely related to JMTV, 9 to Alongshan virus (ALSV), 1 to Xinjiang tick virus 1 (XJTV1), 1 to Yanggou tick virus (YGTV), 5 to Plasmopara viticola lesion associated Jingman-like virus 1 (PVLaJlV1), 16 to Wuhan aphid virus 1 (WHAV1) and 5 to Wuhan aphid virus 2 (WHAV2) strains. We also identified 10 libraries containing a sequence closely related to the known *Ixodes ricinus* jingmenvirus-like segment 1 EVE, and 1 library with a related putative jingmenvirus-like segment 1 EVE from *Ixodes pacificus.* The corresponding 59 libraries are detailed in Table 1, alongside two tick genomic sequence libraries from which we identified novel putative EVEs.

**Table 1:**
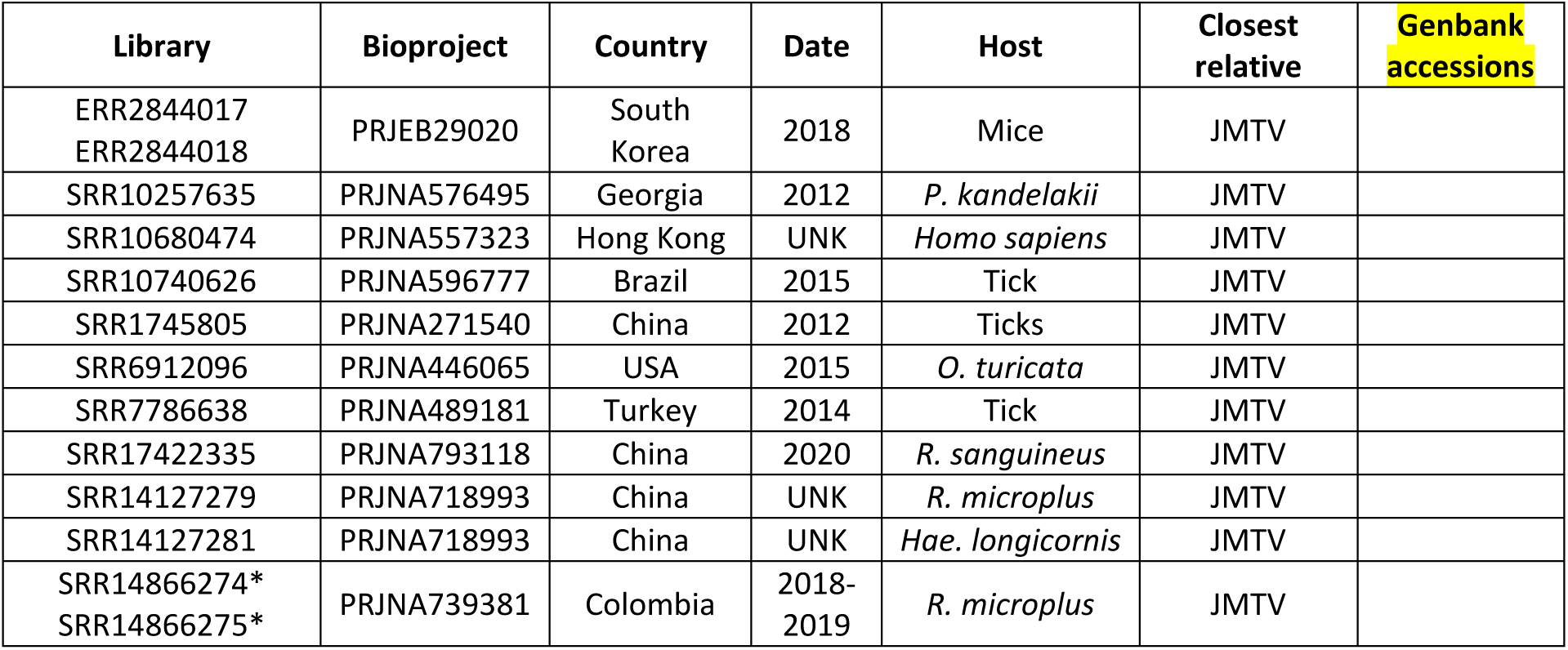
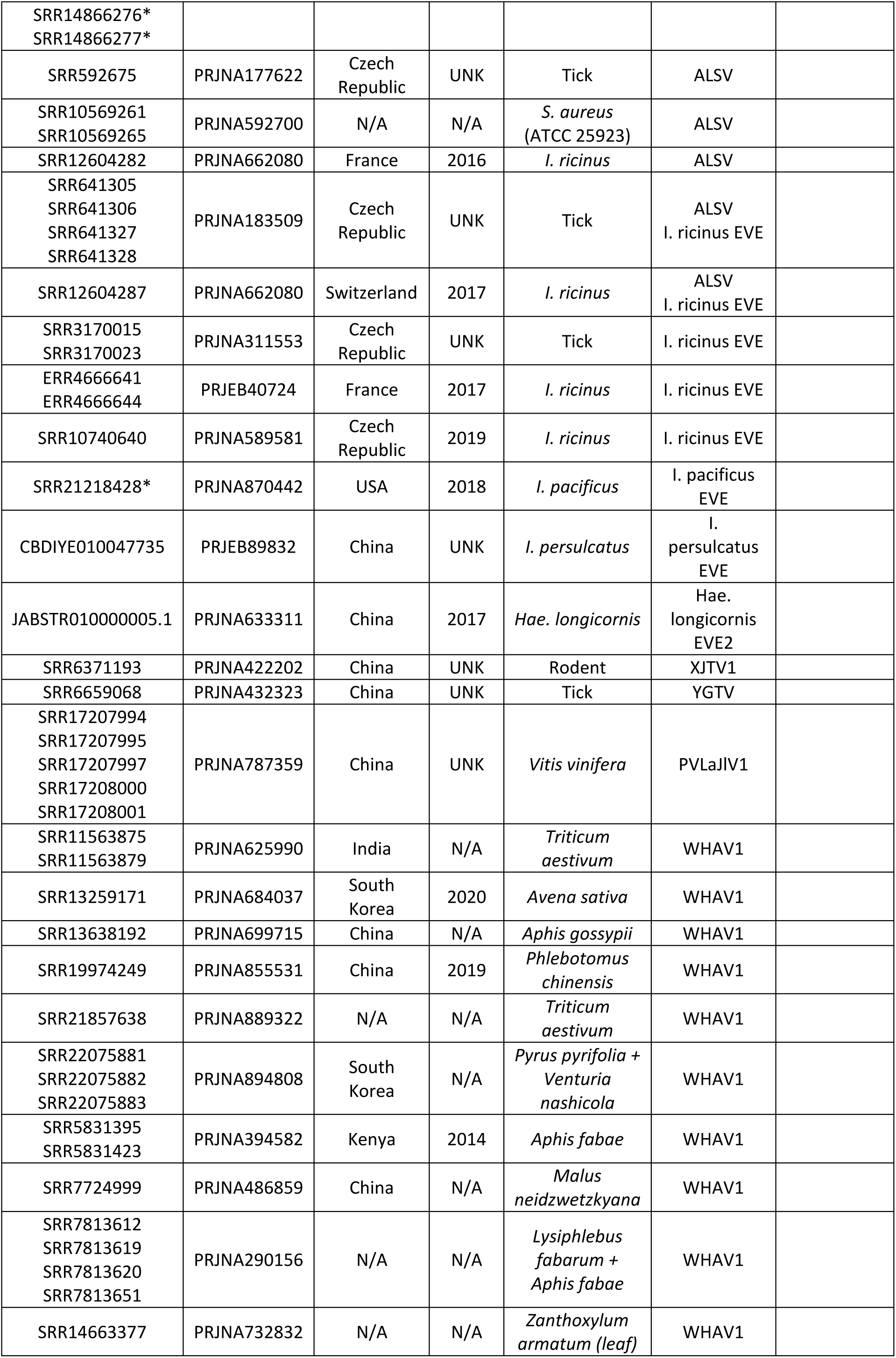
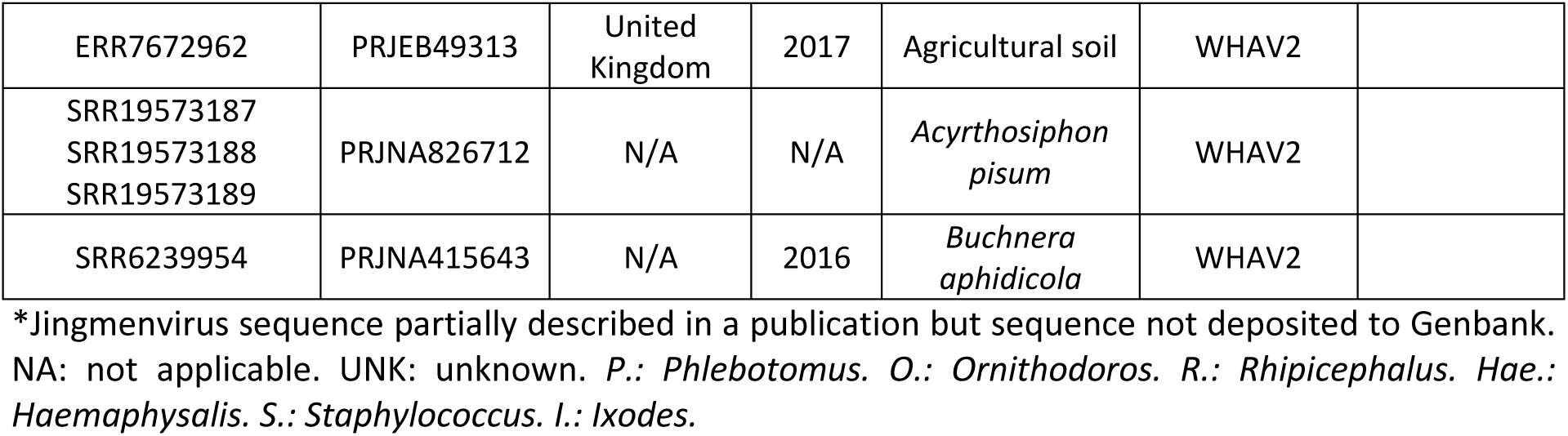
Libraries containing the genomic sequences of new strains of known jingmenviruses or jingmenvirus-derived endogenous viral elements.

| Library | Bioproject | Country | Date | Host | Closest relative | Genbank accessions |
| --- | --- | --- | --- | --- | --- | --- |
| ERR2844017<br>ERR2844018 | PRJEB29020 | South Korea | 2018 | Mice | JMTV |  |
| SRR10257635 | PRJNA576495 | Georgia | 2012 | *P. kandelakii* | JMTV |  |
| SRR10680474 | PRJNA557323 | Hong Kong | UNK | *Homo sapiens* | JMTV |  |
| SRR10740626 | PRJNA596777 | Brazil | 2015 | Tick | JMTV |  |
| SRR1745805 | PRJNA271540 | China | 2012 | Ticks | JMTV |  |
| SRR6912096 | PRJNA446065 | USA | 2015 | *O. turicata* | JMTV |  |
| SRR7786638 | PRJNA489181 | Turkey | 2014 | Tick | JMTV |  |
| SRR17422335 | PRJNA793118 | China | 2020 | *R. sanguineus* | JMTV |  |
| SRR14127279 | PRJNA718993 | China | UNK | *R. microplus* | JMTV |  |
| SRR14127281 | PRJNA718993 | China | UNK | *Hae. longicornis* | JMTV |  |
| SRR14866274*<br>SRR14866275* | PRJNA739381 | Colombia | 2018-2019 | *R. microplus* | JMTV |  |
| SRR14866276*<br>SRR14866277* |  |  |  |  |  |  |
| SRR592675 | PRJNA177622 | Czech Republic | UNK | Tick | ALSV |  |
| SRR10569261<br>SRR10569265 | PRJNA592700 | N/A | N/A | *S. aureus*<br>(ATCC 25923) | ALSV |  |
| SRR12604282 | PRJNA662080 | France | 2016 | *I. ricinus* | ALSV |  |
| SRR641305<br>SRR641306<br>SRR641327<br>SRR641328 | PRJNA183509 | Czech Republic | UNK | Tick | ALSV<br>*I. ricinus* EVE |  |
| SRR12604287 | PRJNA662080 | Switzerland | 2017 | *I. ricinus* | ALSV<br>*I. ricinus* EVE |  |
| SRR3170015<br>SRR3170023 | PRJNA311553 | Czech Republic | UNK | Tick | *I. ricinus* EVE |  |
| ERR4666641<br>ERR4666644 | PRJEB40724 | France | 2017 | *I. ricinus* | *I. ricinus* EVE |  |
| SRR10740640 | PRJNA589581 | Czech Republic | 2019 | *I. ricinus* | *I. ricinus* EVE |  |
| SRR21218428* | PRJNA870442 | USA | 2018 | *I. pacificus* | *I. pacificus*<br>EVE |  |
| CBDIYE010047735 | PRJEB89832 | China | UNK | *I. persulcatus* | *I.*<br>*persulcatus*<br>EVE |  |
| JABSTR010000005.1 | PRJNA633311 | China | 2017 | *Hae. longicornis* | Hae.<br>*longicornis*<br>EVE2 |  |
| SRR6371193 | PRJNA422202 | China | UNK | Rodent | XJTV1 |  |
| SRR6659068 | PRJNA432323 | China | UNK | Tick | YGTV |  |
| SRR17207994<br>SRR17207995<br>SRR17207997<br>SRR17208000<br>SRR17208001 | PRJNA787359 | China | UNK | *Vitis vinifera* | PVLaJIV1 |  |
| SRR11563875<br>SRR11563879 | PRJNA625990 | India | N/A | *Triticum aestivum* | WHAV1 |  |
| SRR13259171 | PRJNA684037 | South Korea | 2020 | *Avena sativa* | WHAV1 |  |
| SRR13638192 | PRJNA699715 | China | N/A | *Aphis gossypii* | WHAV1 |  |
| SRR19974249 | PRJNA855531 | China | 2019 | *Phlebotomus chinensis* | WHAV1 |  |
| SRR21857638 | PRJNA889322 | N/A | N/A | *Triticum aestivum* | WHAV1 |  |
| SRR22075881<br>SRR22075882<br>SRR22075883 | PRJNA894808 | South Korea | N/A | *Pyrus pyrifolia* +<br>*Venturia nashicola* | WHAV1 |  |
| SRR5831395<br>SRR5831423 | PRJNA394582 | Kenya | 2014 | *Aphis fabae* | WHAV1 |  |
| SRR7724999 | PRJNA486859 | China | N/A | *Malus neidzwetzkyana* | WHAV1 |  |
| SRR7813612<br>SRR7813619<br>SRR7813620<br>SRR7813651 | PRJNA290156 | N/A | N/A | *Lysiphlebus fabarum* +<br>*Aphis fabae* | WHAV1 |  |
| SRR14663377 | PRJNA732832 | N/A | N/A | *Zanthoxylum armatum* (leaf) | WHAV1 |  |
| ERR7672962 | PRJEB49313 | United Kingdom | 2017 | Agricultural soil | WHAV2 |  |
| SRR19573187<br>SRR19573188<br>SRR19573189 | PRJNA826712 | N/A | N/A | *Acyrtosiphon pisum* | WHAV2 |  |
| SRR6239954 | PRJNA415643 | N/A | 2016 | *Buchnera aphidicola* | WHAV2 |  |

\*Jingmenvirus sequence partially described in a publication but sequence not deposited to Genbank. NA: not applicable. UNK: unknown. *P.*: *Phlebotomus*. *O.*: *Ornithodoros*. *R.*: *Rhipicephalus*. *Hae.*: *Haemaphysalis*. *S.*: *Staphylococcus*. *I.*: *Ixodes*.

### Building jingmenvirus sequence databases

Using the methods described below, we collated a database of 1,948 published jingmenvirus sequence records across all segments, and added the viral sequences we assembled here for a total of 2,175 sequences (Table 2; Supplementary File 1). 663 were for the segment coding for the RdRp and methyltransferase domains (NSP1), 538 were for the segment coding for the putative glycoprotein (nuORF, VP1, VP1ab, VP4-1 or VP4 and VP5-6), 546 were for the segment coding for the helicase and protease domains (NSP2), 421 were for the segment coding for the putative membrane protein (VP2-3 or VP1-3 and VP2), and 5 were for the fifth segment detected solely in Guaico Culex virus (VP7, thereafter excluded from the analyses). Out of these records 327, 328, 315 and 306 were complete coding sequences of the four segments described above, respectively, out of which 4 x 263 represented putative complete genomes, which we define here as at least 4 segments and complete coding sequences for each segment.

**Table 2:**
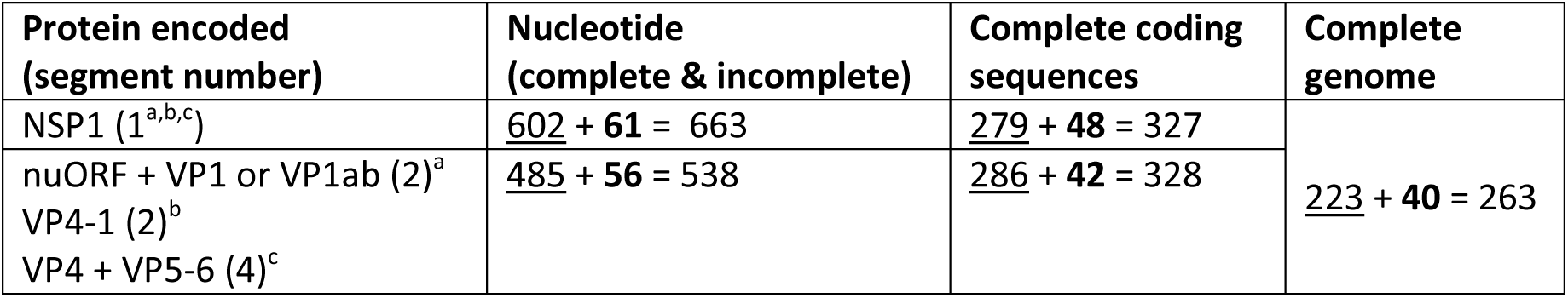
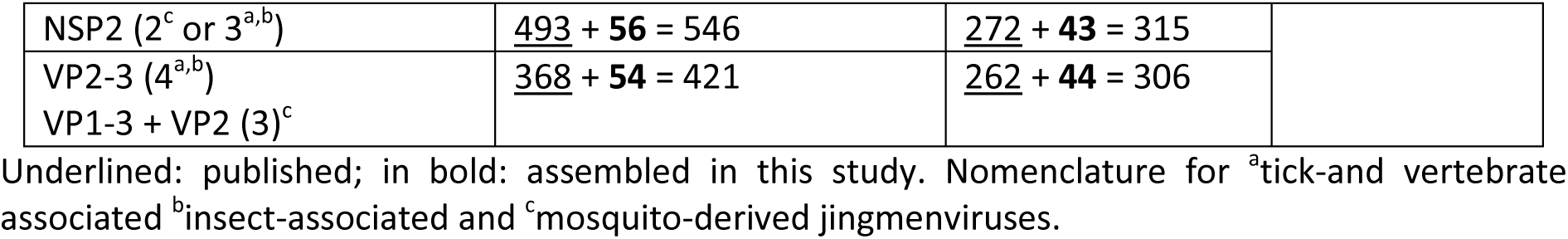
Number of jingmenvirus sequences in the database.

Given the differences in nomenclature between clades, we will thereafter designate segment 1 as the one coding for NSP1, segment 2 as the one coding for nuORF, VP1, VP1ab, VP4-1 or VP4 & VP5-6, segment 3 as the one coding for NSP2 and segment 4 as the one coding for VP2-3 or VP1-3 & VP2.

### Proposing taxonomic classification criteria derived from %ID distribution histograms

The sequences from the 263 complete genomes were used to perform multiple sequence alignments for each segment in nucleotide and amino acid, from which percentage identities (%ID) were inferred. We then derived frequency distribution histograms from the %ID matrices generated. Gaps in this distribution separate groups of viruses that share similar %ID and can denote demarcation criteria to be used to differentiate between virus species (Figure 1). We identified several possible values that could be used as criteria within the distributions obtained. Guided by the phylogenetic trees (Figure 2, Supplementary File 2) derived from the same alignments and by metadata (assigned viral species name, host, location) informing on the biology linked to the sequences, we determined which values were the most relevant to use as criteria to classify sequences into virus species, for all segments, in nucleotide %ID (%IDnt) and %ID amino acid (%IDaa). We aimed at selecting the most parsimonious threshold values that would result in all four segments of a genome being classified the same using the eight %IDnt and %IDaa criteria.

**Figure 1:**
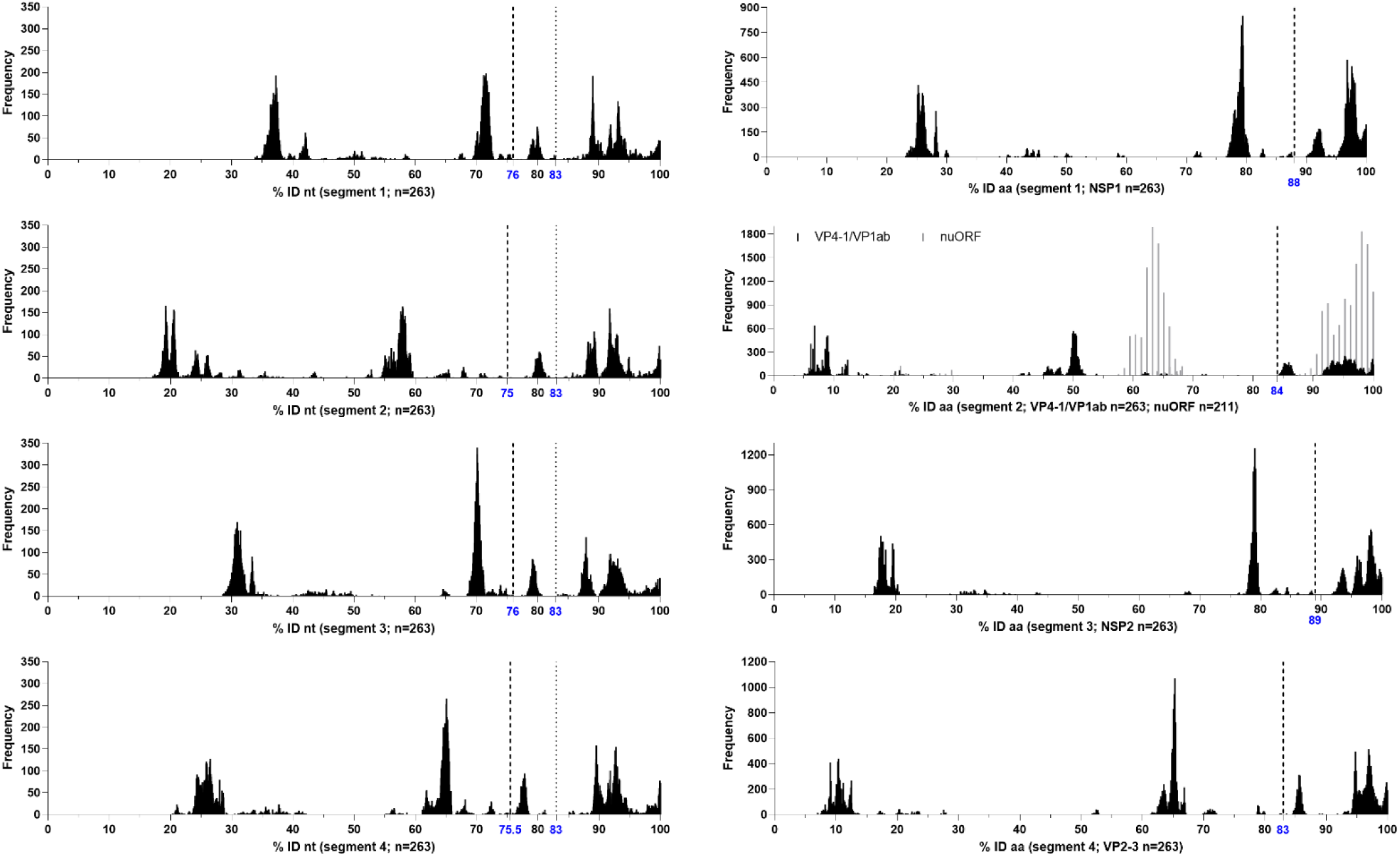
Frequency distribution histograms of percentage identities between jingmenvirus sequences for segments 1 to 4, in nucleotide and amino acid. Dashed vertical lines represent the proposed demarcation criteria for species and dotted vertical lines represent the proposed rule for genotypes. n = 263 for all alignments except nuORF, for which n = 211.

**Figure 2:**
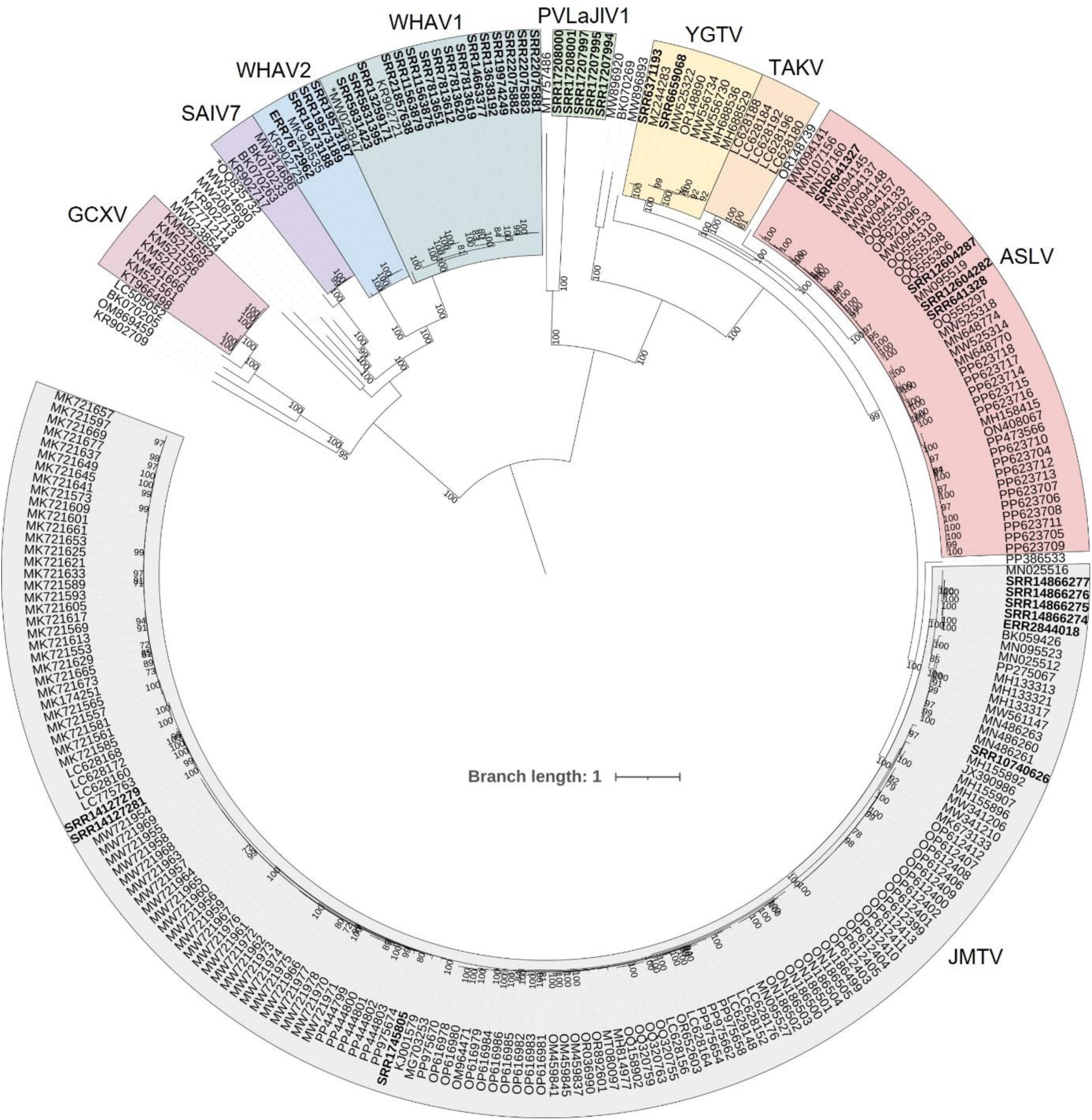
Phylogenetic analysis of segment 1 complete coding sequences (whole genomes only, *i.e.* including at least 4 segments; n=263). The sequences were aligned with MAFFT in Geneious Prime 2024.0.7, the phylogeny was built with IQ-TREE3, using the TIM2+F+I+R5 model and mid-point rooted using Interactive Tree of Life. Bootstrap support values above 70 were included as branch labels. The scale bar represents substitutions per site. The sequences identified by our criteria as part of the same species are highlighted in differently coloured ranges. Sequences that are not in a coloured range represent the only complete genome of their species. The two sequences which we could not conclusively classify using our criteria are preceded by an asterisk *. The sequences assembled in this study are highlighted in bold. JMTV: Jingmen tick virus, ALSV: Alongshan virus, TAKV: Takashi virus, YGTV: Yanggou tick virus, PVLaJlV1: Plasmopara viticola lesion associated Jingman-like virus 1, WHAV1: Wuhan aphid virus 1, WHAV2: Wuhan aphid virus 2,SAIV7: Shuangao insect virus 7, GCXV: Guaico Culex virus.

In nucleotide, we propose that sequences sharing <76%IDnt in segment 1, <75%IDnt in segment 2, <76%IDnt in segment 3 and <75.5%IDnt in segment 4 with their closest relative should be considered a novel species (see Table 3). Taking the segment 1 nucleotide histogram as reference, the other main peaks represent %ID between different species within the tick-associated jingmenvirus clade (60-76%IDnt), between species within the insect-associated jingmenvirus clade (42-60%IDnt) and between species of different clades (< 42%IDnt).

**Table 3:**
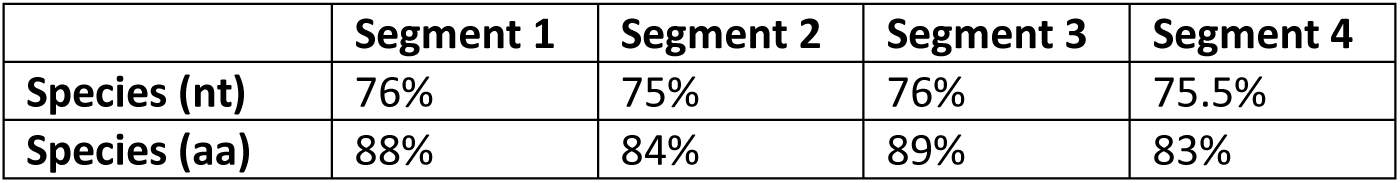
Species demarcation criteria in percentage identity (nucleotide and amino acid)

|  | Segment 1 | Segment 2 | Segment 3 | Segment 4 |
| --- | --- | --- | --- | --- |
| **Species (nt)** | 76% | 75% | 76% | 75.5% |
| **Species (aa)** | 88% | 84% | 89% | 83% |

In amino acid, we propose that sequences sharing <88%IDaa in segment 1, <84%IDaa in segment 2, <89%IDaa in segment 3 and <89%IDaa in segment 4 with their closest relative should be considered a novel species (see Table 3).

We recommend using all criteria when possible, given the segmented and multipartite nature of jingmenviruses, and the associated probability of segment reassortment and/or recombination.

### Applying taxonomic criteria to our database and investigating classification discrepancies

As we recommend the use of the eight %ID criteria (see Table 3), we applied them all to our complete and partial sequences database and found discrepancies between the species our analysis would assign and some sequences published names. We therefore investigated these discrepancies to determine whether our criteria were inaccurate or if they would support re-classification of some published sequences.

#### Grouping sequences published with different names into one species

We found that many sequences associated with various virus names shared %ID way above our proposed thresholds both in nucleotide and amino acid (Table 3) and shared common hosts and/or locations, which would be consistent with these sequences belonging to a single species. In that case, sequences labelled Jingmen tick virus, Mogiana tick virus, Amblyomma virus, Rhipicephalus-associated flavi-like virus, Kindia tick virus, Sichuan tick virus, Guangxi tick virus and Manych virus would all belong to the JMTV species. Following the same rationale, sequences named Harz mountain virus would be included in the ALSV species; sequences labelled Havel Jingmen-like virus and Teltow Canal Jingmen-like virus would be grouped into one species (Teltow Canal jingmen-like virus); sequences labelled Siphonapteran jingmen-related virus isolate OKIAV340 (published and assembled here MW208795, MW208800; new accessions) would be grouped into one species with Wuhan flea virus (WHFV); sequences labelled XJTV1 would be grouped into a single species with YGTV sequences (YGTV), as they fit all nucleotide and amino acid criteria, with only one exception (VP1 81.8%IDaa, thus < 84.0%). We propose to use the name first published for the overall species, or the name linked to the most published segments if published together.

#### Re-classifying to another species

We found sequences sharing >99%IDnt and aa with species that did not correspond to the names they were assigned, so according to our criteria these sequences would be reclassified. Indeed, MN095519-MN095522 and MH678646-MH678648 (labelled JMTV) share >99%ID nt and aa for all segments with strains of ALSV; MG880118-MG880119 (labelled JMTV) share >99%ID with strains of YGTV; OP376740 (labelled YGTV) shares >99%ID with strains of ALSV.

We found that MH155900 is mislabelled as JMTV segment 2, but represents a segment 3 sequence. Then, we found sequences labelled JMTV that shared %ID below all proposed threshold with all described jingmenvirus sequences, which would be classified as new species following our criteria: we propose to designate MN095531-MN095534 as Pteropus lylei jingmenvirus, and MT747997-MT748000 and MT757486 as two strains of Solling histiostoma jingmenvirus. Similarly, OP863303 (labelled JMTV segment 1) shares less than 76%IDnt with JMTV and is more closely related to ALSV, but due to there being only one segment available, we will abstain from classifying it (see below).

#### Re-classifying endogenous viral elements mislabelled as virus

Moreover, we have found sequences labelled as virus that belong to known and previously unidentified EVEs. MT822179, MT822180, OQ162438 (labelled JMTV) and ON933885-ON933950 (labelled Jingmenvirus sp isolate) share >95%IDnt with the previously described *I. ricinus* jingmenvirus-like segment 1 EVE. We found that MZ676705 (labelled ALSV Liaoning; 2.5 kb) and OR114850-OR114911 (labelled Cheeloo jingmen-like virus; up to 2.7 kb) share > 95%IDnt with each other. Since these were the only jingmenvirus-like segment sequences assembled from these samples, we investigated whether they could be integrated in their host genome, based on the existence of the *I. ricinus* jingmenvirus-like segment 1 EVE. In the absence of samples on which to perform molecular analyses, we searched for these sequences using BLAST on the tick’s whole-genome sequences (WGS NCBI database), and found that these sequences indeed corresponded to a jingmenvirus-like segment 2 EVE integration in the *Hae. longicornis* genome. We performed a similar analysis on other single segment sequences for which the host genome had been sequenced and was available and found that OQ716542 (labelled ALSV; 373 bp) is part of another partial segment 2-derived EVE in the *Hae. longicornis* genome, for which we assembled a 1.7kb contig (accession), and which shares 79.0%IDnt with the “Liaoning/Cheeloo” EVE. We found that OP863304 (labelled JMTV; 386 bp), OP244356, OP244397-402, OP244412-14 and PQ043852 (labelled ALSV; 219-332 bp) are part of a jingmenvirus-like segment 4-derived integration in the *I. persulcatus* genome for which we were able assemble a 2.6kb contig (accession). It is important to note that jingmenvirus-derived EVEs can share %ID above our proposed thresholds with jingmenvirus species. For example, the *I. ricinus* jingmenvirus-like segment 1 EVE and *I. pacificus* jingmenvirus-like segment 1 EVE (accession) both share >76%IDnt with ALSV segment 1. However, considering these sequences are not viral RNA but a DNA form integrated in the tick genome, they should be classified as non-viral sequences and therefore not associated to any jingmenvirus species at this stage. The biology of the analysed sequences should be taken into account alongside the %ID criteria when classifying sequences, and the description of novel jingmenvirus species should include the sequences of at least 4 segments to avoid these types of errors.

#### A minority of unclassifiable sequences

For a few sequences, the discrepancies between the published species names and our criteria-derived results were not as straightforward to resolve. Indeed, the eight proposed %ID criteria did not allow the clear classification of three sets of published sequences. First, for MW023847-MW023850, labelled WHAV1, the %IDnt for the four segments are right at the species threshold either just below or just above it, while the four %IDaa criteria would classify the sequences as belonging to a new species. Second, for OQ835732-OQ835735 and BK070268, labelled Jingmenvirus Cameroon (JVC), some criteria would have it classified as belonging to the same species (segment 1, 2-1, 2-2 and 3 nt; segment 2-2 aa) as its closest relative Shuangao insect virus 7 while the other criteria would have it classified as another species (segment 4 nt; segment 1, 2-1, 3 and 4 aa). Third, the sequence OP863303 (labelled JMTV segment 1) would belong to a novel jingmenvirus species based on the %IDnt criterion, but its NSP1 ORF shares 100%IDaa with an ALSV strain (QHW06950). We intended to find other segments in the associated libraries to help with classification, but we did not find any reads mapping to this sequence so it is unclear how the authors assembled it in the first place (14). Considering there was a single segment published, we searched for but found no evidence that this sequence could be integrated in the *I. persulcatus* genome with BLAST wgs.

The proposed criteria allowed to identify these sequences as outliers but with the current lack of published data on the biology of these viruses, it is not clear yet how they should be classified. Characterising the viruses in question would allow to definitively conclude on the matter.

Considering that the ten unclassifiable sequences described in the previous paragraph represent a small minority within the large 2180 sequences database, we are satisfied with the performance of the eight proposed criteria, with the addition of the recommendation to have at least 4 segment sequences when describing a new species.

### Criteria with >99.6% accuracy performance

We endeavoured to confirm the performance of the proposed eight criteria by measuring their accuracy on the complete coding sequences database. In order to do so, we compared the %IDnt and %IDaa of each sequence with every other sequence in the database (not only their closest relative) and counted how many did not fit the criteria properly: for example, in the segment 1 %IDnt matrix, how many %ID were below 76% in members of the same assigned viral species and how many were above between members of different species. We then calculated the percentage accuracy of each criterion by comparing that number to the total number of entries in each %ID matrix. We classified the sequences according to our criteria, and chose a reference for the outlier sequences (MW023847-MW323850 sequences considered as WHAV1; OQ835732-OQ835735 and BK070268 as its own species). With these species assignment, the accuracy of the method is >99.6% for all criteria (Table 4).

**Table 4:**
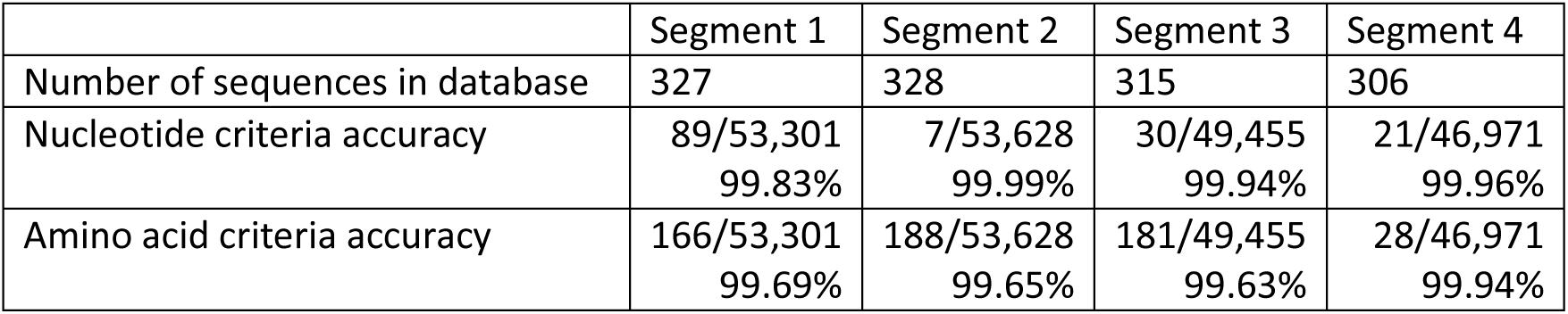
Classification criteria accuracy. Number of %ID value outside of the proposed criteria in each %ID matrix, based on the CCDS database; and corresponding percentage accuracy for each criterion.

### Identifying several jingmenvirus species with two genotypes/lineages

Considering that we and others have described numerous and sometimes divergent strains for a few jingmenvirus species, we also propose a rule for genotype/lineage identification with a threshold at 83%IDnt for all segments. While this rule is out of the scope of taxonomical considerations, it will benefit the jingmenvirus research community by clarifying the spatial and temporal distribution as well as host association of divergent strains of the same viral species. We opted to number putative genotypes (I and II) based on the chronology of their publication: earlier sequences being assigned to genotype I.

Following this rule, we found that Jingmen tick virus could be separated in two genotypes. JMTV genotype I would include complete and partial sequences from Asia (China, Japan, Laos, Hong Kong), Africa (Kenya, Uganda, Guinea, Cameroon), Europe (Georgia, Turkey, Russia) and the Americas (Brazil, USA). The JMTV genotype I sequences would originate from a wide range of hosts: arthropods such as ticks (genera *Amblyomma, Dermacentor, Haemaphysalis, Hyalomma, Ixodes* and *Rhipicephalus* as well as the soft tick *Ornithodoros turicata*), mosquitoes (*Armigeres sp.*), sand flies (*Phlebotomus kandelakii*) and vertebrates such as rodents (genera *Apodemus, Cricetulus, Meriones, Microtus, Mus, Rattus* and *Rhombomys*), bats (genera *Eptesicus, Miniopterus, Myotis, Nyctalus, Pipistrellus* and *Rhinolophus*), cattle (*Bubalus bubalis* and *Bos taurus*), primates (*Piliocolobus rufomitratus* and humans) and reptiles (*Stigmochelys pardalis*). JMTV genotype II would include complete and partial sequences from Europe (Kosovo, Romania, Turkey, Italy, France, Russia), the Americas (Trinidad and Tobago, Guadeloupe/Martinique, Colombia, Mexico) and Asia (South Korea; although it is not clear whether the sequence originated from laboratory or environmental sample).

The sequences grouped in this JMTV genotype would originate from different host types: tick species such as *R. microplus, R. bursa, R. sanguineus, R. turanicus, H. marginatum, I. simplex* and *Am. variegatum,* other arthropods such as *Ae. albopictus* from a laboratory colony and *Ochlerotatus caspius*, human patients and mice intestine of unknown species and origin (laboratory or environmental). Strikingly, the segments of a given JMTV strain all fall in the same genotype. Moreover, we found only one study with detections of both JMTV genotypes in tick-derived samples from the same country (Turkey) with 2 out of 13 strains partially sequenced over segment 3 belonging to JMTV genotype II and the rest belonging to JMTV genotype I (15). Additionally, two separate studies have found partial segment 2 sequence of both JMTV genotypes in Russia (MN218697 and MN218698 JMTV genotype I, PQ310748 JMTV genotype II). These are the only two countries to date with both JMTV genotypes recorded.

While seven out of eight %ID criteria would group YGTV and XJTV1 in a single species (YGTV), they share %IDnt that would classify them as two putative genotypes, with particularly divergent VP1 sequences, which could have evolved under different environmental pressures. The sequences formerly identified as YGTV would form genotype I and the sequences formerly identified as XJTV1 would form genotype II. YGTV genotype I would have been detected in *D. nutalli, D. marginatus, D. reticulatus, D. silvarum* and *I. persulcatus* ticks collected from Russia and China. YGTV genotype II would have been detected in rodents and ticks collected from China.

Similarly, while our eight %ID criteria would group WHFV and OKIAV340 into a single species, their genetic distance would classify them as two distinct genotypes. The sequences formerly identified as WHFV would form genotype I and the sequences formerly identified as OKIAV340 would form genotype II. WHFV genotype I would have been detected in *Ctenocephalides felis* from China, and WHFV genotype II would have been detected in the same host collected from the USA.

While all GCXV isolates are clearly part of the same species, one strain (KM521571-KM521574) has divergent segments 2, 3 and 4 that would be classified as a different genotype, which would suggest this isolate should be characterised to determine whether these nucleotide differences in 3/5 segments result in phenotypical alterations.

### Phylogenetic analysis is in accordance with %ID criteria

We built phylogenetic trees for the sequences from complete genomes using the complete coding sequence alignments described above (Figure 2, Supplementary File 2) and to visualise and validate the relevance of the proposed demarcation criteria. It is clear in this representation of evolutionary distance that the sequences classified as the same species are closely related and that the sequences classified as separate species are evolutionarily distant. We propose to classify 25 jingmenvirus species corresponding to the genomes with at least 4 complete segments which are sufficiently different from each other and from their closest relatives, according to our proposed criteria (Supplementary File 1). Among these 25 proposed species, 9 species include several complete genomes (see coloured ranges in Figure 2 and Supplementary File 2).

The unclassified sequences included in these analyses (preceded by an asterisk on the figure) were indeed in intermediary positions on the tree, in the same clade as their closest relative but a bit more distant than the other members of the species, and this distant varied between segments and between nucleotide and amino acid analyses (Figure 2, Supplementary File 2).

The whole genome sequences included in these analyses for which we suggest grouping in two genotypes (JMTV and YGTV) have two clear clusters in their clade, corresponding to our genotype proposals.

## Discussion

While this manuscript was in its final stages of submission, a proposition to reorganise the taxonomy of the *Flaviviridae* was published by the corresponding ICTV Study Group (16). In that study, the authors propose to organise the current *Flaviviridae* in 3 families (*Flaviviridae, Pestiviridae, Hepaciviridae*), which would include 14 genera. Two of the new genera would include segmented flavi-like virus sequences, the proposed *Jingvirus* genus with Jingmen tick virus as a single recognised species and the *Guaicovirus* genus with Guaico Culex virus as a single recognised species, both part of the revised *Flaviviridae* family. This show that there is indeed a great need to clarify the organisation of flavi-like sequences and their taxonomy, and the species-level criteria we propose for jingmenviruses (or segmented flavi-like viruses) complement these propositions perfectly.

First, we report the detection of new sequences from several jingmenvirus strains, extending the known spatial and host range for these virus species. These new sequences were assembled from publicly available raw sequencing data published by others. The use of these results demonstrates the importance of open access data publication, with accurate associated metadata. By combining these newly assembled sequences with the existing ones, we collated a database that reached critical mass for addressing the taxonomic organisation rules of jingmenviruses. Indeed, we have determined eight %ID criteria (Table 3), for four segments in nucleotide and amino acid, which we propose could be used to help taxonomically classify jingmenvirus sequences. We propose that four complete segments sequences sharing less than 75-76%IDnt and/or 83-89%IDaa (depending on the segment, see Table 3) with their closest relative could be considered as a novel jingmenvirus species. The cut-off values determined in this study are lower than those applied to members of the *Flavivirus* genus (84%IDnt over the whole genome) (17). This suggests that while these groups of viruses seem related, as they share genetic similarity over the segments coding for non-structural proteins, jingmenviruses have a wider diversity than flaviviruses.

We are proposing %ID criteria that enabled us to (i) clearly identify currently published jingmenvirus species; (ii) rationally propose to group sequences labelled with different species names into a single species; (iii) propose to re-classify sequences labelled with the name of another viral species than their closest relative; (iv) identify EVEs from tick genomes; and (v) single out sequences with peculiar profiles which would benefit from *in vitro* characterisation. The existence of criteria brings some level of order to a group of viruses unclassified by the ICTV, and will help avoid labelling errors being perpetuated in various publications.

Beyond taxonomic classification, a rule for genotypes/lineages has enabled us to show that the widely accepted notion that jingmenvirus sequences group together based on their location might not be completely accurate (15,18). Indeed, we found that both JMTV genotypes have been detected in Turkey and Russia, from the same location and the same tick species, and that both YGTV genotypes have been detected in China (15,19). Moreover, with JMTV sequences separated into two groups, we have been able to look for reassortments between the two genotypes and found no evidence of this phenomenon. This is not unexpected, considering that out of all studies reporting the detection of JMTV, only one has found both genotypes in the same location and tick species (15). It therefore seems that interactions between the two JMTV genotypes, while possible, are uncommon. Similarly, the two WHFV putative genotypes were detected from the same host (*Ctenocephalides felis*) but from different locations (China and USA), so there is no evidence that they share an ecological niche which would facilitate reassortment. The two YGTV putative genotypes have been found in China, but it is unclear whether they share a host, as genotype II was found in samples with metadata listing “tick” and “rodent” hosts, and genotype I was found only in ticks, of the *Dermacentor* and *Ixodes* genera or unspecified genus. What is clear is that to date, no reassortment was observed between genotypes of any of these jingmenvirus species, irrespective of whether they share ecological niches or not (20).

We believe that proposing and using the eight %ID criteria (four segments, nt and aa) is essential, in particular since jingmenviruses are segmented and thought to be multipartite. Indeed, when investigating discrepancies between published names and our analysis, we identified outlier sequences with different genetic distance profiles which could be linked to mechanisms that drove their evolution. For example, sequences closely related in amino acid and more divergent in nucleotide could be the product of constraints from different hosts shaping the nucleotide composition and codon usage, while the constraints on protein function and therefore amino acid sequence remain. Or, finding a discrepancy of genetic distance between structural *vs* non-structural genes could highlight evolutionary pressures acting only on selected aspect of the viral life cycle. Moreover, jingmenviruses are multisegmented and multipartite, meaning that they would encapsidate less fragments than the complete genome and that multiple particles would be required for infection (21–23). With this type of organisation, it is conceivable that one divergent segment could become part of a jingmenvirus infectious unit through co-infection or intra-host evolution (4). Identifying such profiles is key to understanding the evolutionary mechanisms, life cycle and biology of jingmenviruses.

We have uncovered multiple instances of single segment sequences being labelled with jingmenvirus species names, when in reality they were integrated in their tick host genome. When using only the proposed %ID criteria, these sequences could indeed have passed as strains of existing species, but uncovering and taking into account their biological origin was key to classifying these sequences correctly. We therefore recommend to obtain at least 4 segments to consider that the new sequences indeed form a viral genome, as the currently recognised minimal number of segments for an infectious unit is four (4,21,24). We propose that novel jingmenvirus-related sequences with less than four segments should not be labelled as virus species but rather as virus-related sequences.

Another reason to look for more than one jingmenvirus-related segment in sequencing data before labelling or classifying it would be to find out whether the sequence could be a previously undetected part of an existing genome, as we found in a previous study. Indeed, jingmenviruses can have up to at least 6 segments, with pairs of segments coding for homologous structural proteins (4). In particular, we uncovered a previously unknown fifth segment for SAIV7 and JVC, homologous to their respective segment 2. The fluid genomic organisation of jingmenviruses could complicate the classification of jingmenviruses, considering that in all documented cases, the two homologous segments from the same species are sufficiently genetically distant to be considered as different species, when compared. This should be taken into account, and particular attention to assembling as complete a genome as possible should be paid when describing new jingmenvirus sequences. A multisegmented multipartite organisation facilitating segment duplication, loss or acquisition, intra-and inter-host evolution or reassortment complicates the definition of a jingmenvirus species.

The criteria we propose and use here are based on current knowledge of jingmenvirus genomic organisation and biology. They are susceptible to change when further characterisation and discoveries lead to a better understanding of these viruses. A great example of re-classification based on further knowledge are jiviviruses, which were originally thought to include virga-like and jingmen-like segments coding for non-structural proteins, and segments coding for putative structural proteins of unknown origin (25,26). With the significant increase of sequencing data in the era of viromics, these considerations have since been revised, and jivivirus non-structural proteins have been shown to be more closely related to the monopartite families *Potyviridae* and *Flaviviridae* (27). The existence of such viral genomes borrowing segments or domains from different viral families are the perfect example of why multiple criteria should be met to classify new sequences as a novel jingmenvirus species.

In conclusion, based on the current state-of-the-art, we recommend to only name novel jingmenvirus sequences as novel species when at least four segments have been sequenced, and they share <76%IDnt and <88%IDaa in segment 1, <75%IDnt and <84%IDaa in segment 2, <76%IDnt and <89%IDaa in segment 3, <75.5%IDnt and <83%IDaa in segment 4 with their closest relative.

## Material and methods

### Assembling new jingmenvirus strains from publicly available sequencing libraries

We used the online Serratus.io platform to find novel jingmenvirus sequences in publicly available raw sequencing data from the sequence reads archive (SRA) (28). We started by searching for novel Jingmen tick virus (JMTV) related sequences. We used the Serratus NT Search (option Genbank) for JMTV strain SY84 segment 1 (NC_024113.1), selecting only libraries that matched with a score between 3 and 100, and with a percentage identity above 75%IDnt, hoping to uncover libraries with JMTV sequences as well as sequences of other jingmenvirus species in the tick- and vertebrate-associated clade. We obtained 92 libraries that matched these criteria.

We then broadened the scope of our search and used the Serratus RdRp Search (option Family) for *Flaviviridae*, with a score criteria between 50 and 100, and a percentage identity above 90%IDnt. We obtained 9,615 libraries sorted in subfamilies numbered by the Serratus tool. We identified the jingmenvirus-related subfamilies: Flaviviridae-3 (tick-associated jingmenviruses) and Flaviviridae-28 (PVLaJlV1) and the 63 associated libraries (56 for Flaviviridae-3, 7 for Flaviviridae-28) (26,27,29).

Finally, we aimed at finding sequences related to jingmenviruses in the insect-associated clade. We therefore selected the Serratus RdRp Search and screened for libraries in the relevant “Family”, entitled Unclassified-899. With the filtering criteria used (percentage identity > 90%IDnt and score > 50) we identified 43 libraries.

We then selected only the libraries which were not associated with publications on jingmenviruses, eliminated duplicate records generated by our different screens and assembled jingmenvirus-related sequences from 59 libraries. We used NCBI BLAST with its direct access to the SRA to obtain reads mapping to reference sequences, with the Megablast algorithm optimised for highly similar sequences with maximum target sequences set to the highest setting (5000) and assembled these reads using Geneious Prime 2024.0.7. The following reference sequences were used: JMTV KJ001579-KJ001582; ALSV MH158415-MH158418; YGTV MH688529-MH688532; XJTV1 MZ244282-MZ244285; PVLaJlV1 MN551114, MN551116, BK061346, BK061347; WHAV1 KR902721-KR902724; WHAV2 KR902725-KR902728 and Ixodes ricinus NSP1 EVE OP068155.

We included partial sequences in the database when the most represented segment had been assembled with at least 25 reads.

### Database of published jingmenvirus sequences

We endeavoured to build a database for all published jingmenvirus sequences to be used in taxonomic analysis in addition to the newly assembled sequences. In order to do so, we compiled all sequences obtained with a search on NCBI Genbank with the following keywords: jingmenvirus, jingmen-related, jingmen-like, segmented flavivirus, segmented flavi-like, flavi-like segmented virus, flavivirus-like, flavi-like, Jingmen tick virus, Mogiana tick virus, Kindia tick virus, Guangxi tick virus, Manych virus, Sichuan tick virus, Alongshan virus, Yanggou tick virus, Xinjiang tick virus, Takachi virus, Guangdong jingmen-like virus, Hainan jingmen-like virus, Pteropus lylei jingmenvirus, Peromyscus leucopus jingmenvirus, Guaico Culex virus. We then searched NCBI Pubmed with the same keyword list and compiled all published sequences related to the articles uncovered by that search. We also screened all articles citing Qin *et al.* (30), the first description of a jingmenvirus, and compiled all jingmenvirus sequences from these articles. Finally, we used NCBI BLASTx and searched for sequences related to all jingmenvirus species identified until then, adding to our database the sequences listed in the results that slipped through the screening process.

Our database was last updated with newly published sequences on April 11^th^ 2025.

### Percentage identity frequency distribution histograms

Nucleotide and protein sequences were aligned using MAFFT v7.490 (31) with the L-INS-i algorithm, which is recommended for sequences with conserved domains and variable regions. The L-INS-i strategy uses iterative refinement with local pairwise alignment information and is particularly suitable for datasets with multiple distinct sequence regions.

We selected only records with complete genomes (at least 4 complete coding sequences) and aligned their coding nucleotide sequences as well as their amino acid translation for all segments, except for records of Guaico Culex virus (GCXV) segment 5 since they have no known homologs. Given the differences in nomenclature between clades, we will thereafter designate segment 1 as the one coding for NSP1, segment 2 as the one coding for nuORF, VP1, VP1ab, VP4-1 or VP4 & VP5-6, segment 3 as the one coding for NSP2 and segment 4 as the one coding for VP2-3 or VP1-3 & VP2. We exported the percentage identity matrices for all alignments and used the FREQUENCY function of Excel to generate a table of frequencies, with percentage identity values rounded to the hundredth as bins. The frequencies were plotted as distribution histograms with GraphPad Prism 9. Frequencies that are not represented in the original matrices result in gaps in the distribution, which indicates a separation between groups of sequences, and can be used as criteria for taxonomic classification (11,12).

### Phylogenetic analysis

Best-fit evolutionary models for both nucleotide and protein alignments were determined using ModelFinder (32) as implemented in IQ-TREE 3.0.1. Model selection was performed based on the Bayesian Information Criterion, which balances model fit and complexity to avoid overfitting (Table 5). For nucleotide alignments, the selected models ranged from TIM2+R4 to TIM2+F+I+R5, indicating varying levels of rate heterogeneity across different genomic segments. For protein alignments, the selected models included LG+I+R4, LG+I+R5 and JTT+I+R4, reflecting the diverse evolutionary constraints acting on different protein-coding regions.

**Table 5:**
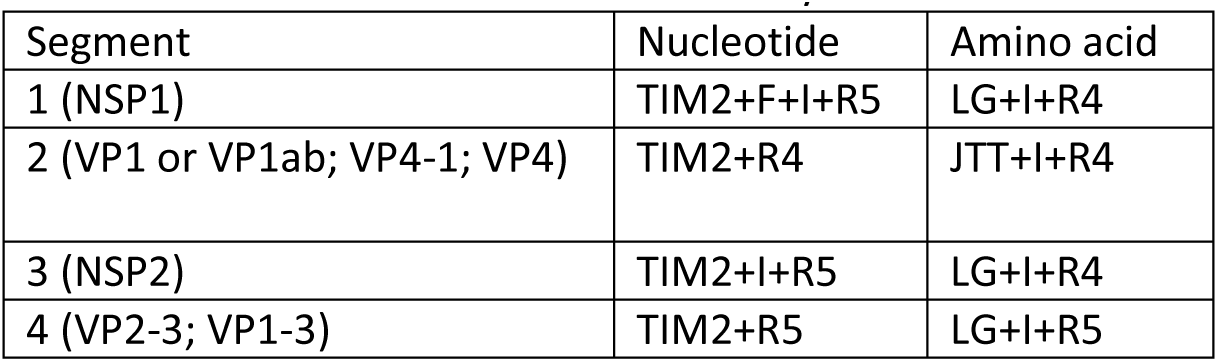
Models selected based on the Bayesian Information Criterion for phylogenetic analyses.

Phylogenetic trees were constructed using IQ-TREE multicore version 3.0.1 for Windows (33) with the best-fit models as determined by ModelFinder. Branch support was assessed through 1000 ultra-fast bootstrap replicates (34) and 1000 SH-like approximate likelihood ratio test replicates (35). The command used for phylogenetic analysis was: iqtree3 -s [alignment_file] --alrt 1000 -B 1000. The trees were visualised, mid-point rooted and formatted using the online platform Interactive Tree of Life, version 7.2.2 (36).

## Data availability

The sequences assembled in this study were deposited on NCBI Genbank as third party annotations and are accessible under the following accession numbers: xxxxxxxx-xxxxxxxx and xxxxxxxx-xxxxxxxx. The details for all sequences in the database can be found in Supplementary File 1.

## Supplementary Files legend

Supplementary File 1: Sequences in jingmenvirus database, with metadata and proposed or tentative classification.

Supplementary File 2: Phylogenetic analyses of segments 1, 2, 3 and 4 complete coding nucleotide and amino acid sequences (whole genomes only, i.e. including at least 4 segments; n=263). The sequences were aligned with MAFFT in Geneious Prime 2024.0.7, the phylogeny was built with IQ-TREE3, using the model described in Table 5 respectively and mid-point rooted using Interactive Tree of Life. Bootstrap support values above 70 were included as branch labels. The scale bar represents substitutions per site. The sequences identified by our criteria as part of the same species are highlighted in differently coloured ranges. Sequences that are not in a coloured range represent the only complete genome of their species. The two sequences which we could not conclusively classify using our criteria are preceded by an asterisk *. The sequences assembled in this study are highlighted in bold. JMTV: Jingmen tick virus, ALSV: Alongshan virus, TAKV: Takashi virus, YGTV: Yanggou tick virus, PVLaJlV1: Plasmopara viticola lesion associated Jingman-like virus 1, WHAV1: Wuhan aphid virus 1, WHAV2: Wuhan aphid virus 2,SAIV7: Shuangao insect virus 7, GCXV: Guaico Culex virus.

## Acknowledgments

This work was partly supported by the European Viral Outbreak Response Alliance EVORA project under HORIZON EUROPE framework (Grant agreement ID: 10113195). The funders had no role in study design, data collection and interpretation, or the decision to submit the work for publication.

